# New Perspective on GWAS: East Asian Populations from the Viewpoint of Selection Pressure and Linear Algebra with AI

**DOI:** 10.1101/2022.05.27.493710

**Authors:** Masayuki Kanazawa

## Abstract

Genome Wide Association Studies (GWAS) are useful for comparing the characteristics of different human population groups. However, genomes can change rapidly over time when there is a strong selection pressure, such as a pandemic. The genetic information related to the immune system is thought to be very sensitive to such diseases. Therefore, it may be necessary to conduct not only the standard whole-genome GWAS but also a more detailed, chromosome-focused GWAS.

In this study, we compared chromosomes of immune system genes to those that are not thought to be related to the immune system, and analyzed GWAS results for SNPs in each chromosome to examine the differences. In order to keep the sample conditions as identical as possible, we limited the comparisons and the analyses to a few groups for which population movements were easy to interpret, and we also made sure the sample sizes were as close as possible. We selected a population of 403 East Asian people, consisting of 104 Japanese people in Tokyo (JPT), 103 Han Chinese people in Beijing (CHB) and 105 in southern China (CHS), and 91 Korean people (KOR). PCA and Manhattan plot were used to analyze and compare the results.

Japanese, Chinese, and Korean populations formed distinctly different groups, with major differences observed. Validity of PCA and Manhattan plot was also discussed using Mahalanobis distance and AI.

## 1. Introduction

### 1.1 Background

Rapid advances in DNA sequencing technologies in recent years have led the International Human Genome Sequencing Consortium to announce the successful completion of the decoding of the human genome in 2003 [1]. Subsequently, a global whole-genome sequencing project (the 1000 Genomes Project) was initiated in 2008 with the goal of characterizing the genetic diversity of the world’s population. As a result, genome information on more than 2,500 individuals worldwide is presently available as open data, including Japanese and Chinese individuals in East Asian populations [2-3]. More data are now available from other populations, including Koreans and Mongolians [4-5], which not been covered by the 1000 Genomes Project.

These data are being analyzed mainly by linear algebraic methods, such as Genome Wide Association Studies (GWAS), and are beginning to reveal genomic characteristics in European populations and details of human inter- and intra-regional migrations from the past to the present [7]. Recently, studies have been conducted not only in Europe but also in East Asian populations such as Japanese, Chinese, and Korean populations [8-13].

### 1.2 Issues of Previous Studies

Most of the previous GWAS that compared human groups have focused on whole genomes or on genes associated with specific infectious diseases. However, not many studies have analyzed these associations at the chromosomal level, even though it is chromosomes where the genetic recombination is actually occurring. There are also several linear algebraic issues.

First, GWAS on whole genomes do not usually assume that genes change significantly over time due to a strong selection pressure; however, the genomes of the immune system, such as Human Leukocyte Antigen (HLA) system, are expected to change significantly in response to pandemics. Therefore, when the results of Principal Component Analysis (PCA) on Single Nucleotide Polymorphisms (SNPs) obtained by GWAS are used to estimate group associations back in time, it is not always guaranteed that the data are consistent from the past to the present. This may reduce the accuracy of the analysis results.

Second, in the previous studies, the volumes of sample sizes of the groups to be compared were often different; PCA on SNPs, which is often employed in GWAS, has the property of determining the scale (principal components) so as to ensure that the total variance of all samples is maximized. Therefore, the principal components that serve as the scale for comparison of all groups will be optimized for the group with the largest sample size, which may not necessarily guarantee total optimization. Thus, even if the same groups are compared, it is not uncommon among studies with different sample sizes to yield different results.

### 1.3 Aim of This Study

To address the above issues, this study adopted the following alternative research approaches. First, comparisons will be limited as much as possible to only a few groups for which population movement can be easily interpreted. Second, chromosomes with immune system genomes that are likely to change frequently will be selected, along with chromosomes that are not, and then the results of PCA of SNPs by GWAS for each chromosome will be compared.

Specifically, in the East Asian populations of Japanese, Chinese, and Korean subjects that are currently available as open data, chromosomes that contain genes closely related to infectious diseases and other chromosomes will be analyzed by PCA and Manhattan plot.

## 2. Methods

### 2.1 Genome Data Used

Genome data from the aforementioned 1000 Genomes Project were used; they are publicly available as open data and include people in Tokyo, Japan (104 Japanese; JPT), Beijing, China (103 Han Chinese; CHB), and southern China (105 Han Chinese; CHS) as East Asian populations [2]. For Koreans, we used data from the KPGP that is currently available as open data (91 Koreans; KOR) [14]. The number of people in each group is about the same (around 100), and the number of males and females is also about the same, which is expected to reduce PCA bias. Since these data are open data, anyone can use them freely.

The following advantages can also be expected from the data used in this study.

1. The geographical proximity and small number of groups make it relatively easy to verify the results of the analysis.
2. China has a long history, and events that may affect the genome (such as population movements, wars, and pandemics) are available in historical books and other written records, and can be compared with GWAS results.
3. Population sizes of these countries are relatively large (Japan 126m, China 1,380m, Korea 52m) and group members are stable.

### 2.2 Target Chromosomes

Based on the aforementioned reasons, the following chromosomes were selected for analysis, referring to the genome map created by the Japanese Ministry of Education, Culture, Sports, Science and Technology [15].

1. Chromosome 1, which is a common chromosome and is assumed to contain few genes related to the immune system.
2. Chromosome 6, which contains HLA genes, and is the center of the human immune system.
3. Chromosome 9, which contains the gene for ABO blood group, and has been studied in relation to many infectious diseases including COVID-19 [16-17].
4. Chromosome 12, which contains ALDH2 (Human Aldehyde Dehydrogenase 2), the “alcohol-sensitivity”; information encoded on this chromosome tends to be similar among people in Japan, southern China, and Korea, while also being considered to be distinct from other human groups [18].

### 2.3 Analysis Methods

PCA was conducted for the above chromosomes, and the first and the second principal components were analyzed to calculate the Mahalanobis distances from JPT to CHB, CHS, and KOR. Similarly, analysis using Manhattan plot was conducted for each chromosome comparing the Japanese population to the others. The software used was plink v1.9 and 2.0 [19] for PCA and R v4.1.3 [20] for Manhattan plot. The alpha level was set at 0.05 and Bonferroni’s correction was used.

Chromosome 6, which contains the genes of HLA, a major immune system, was divided into the first half, in which the genes of HLA are present, and the second half, in which they are absent. PCA was performed on each portion. The numbers of SNPs in the first half and the second half were set equal.

## 3. Results

### 3.1 Chromosome 1

In the first and second principal components, where the differences were largest, the Japanese, Korean, and Chinese populations were clearly separated. The groups of Beijing (CHB) and southern China (CHS) also had their own characteristics, but as a whole they constituted one group (Fig. 1). Manhattan plot showed no noticeable difference (Fig. 2).

**Fig. 1.**
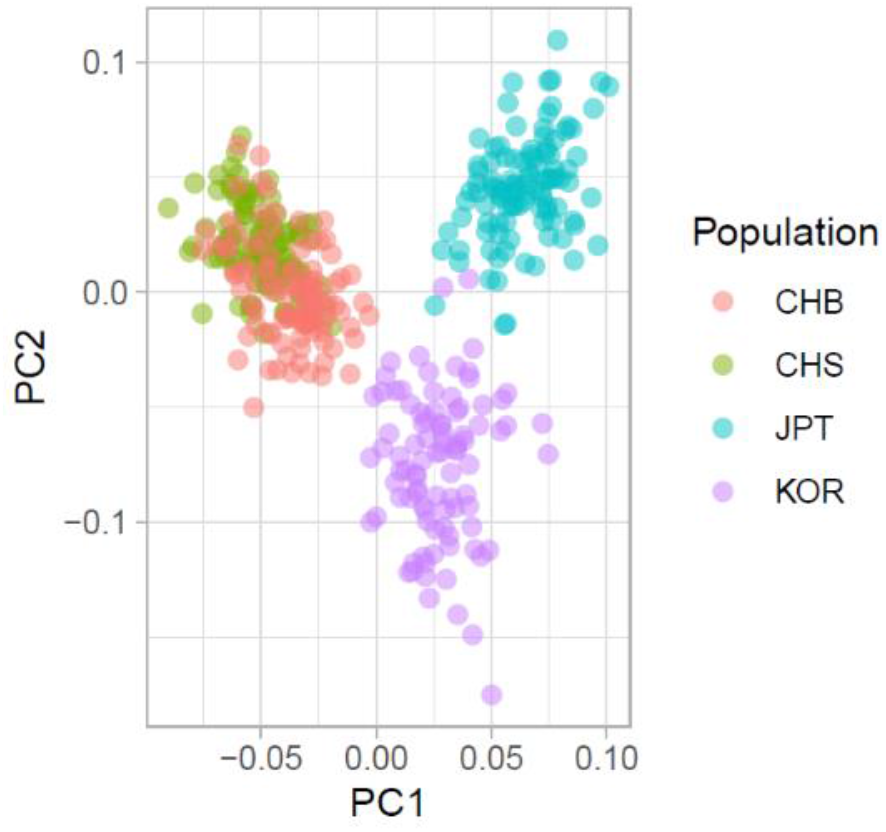
PCA Result of Chromosome 1.

**Fig. 2.**
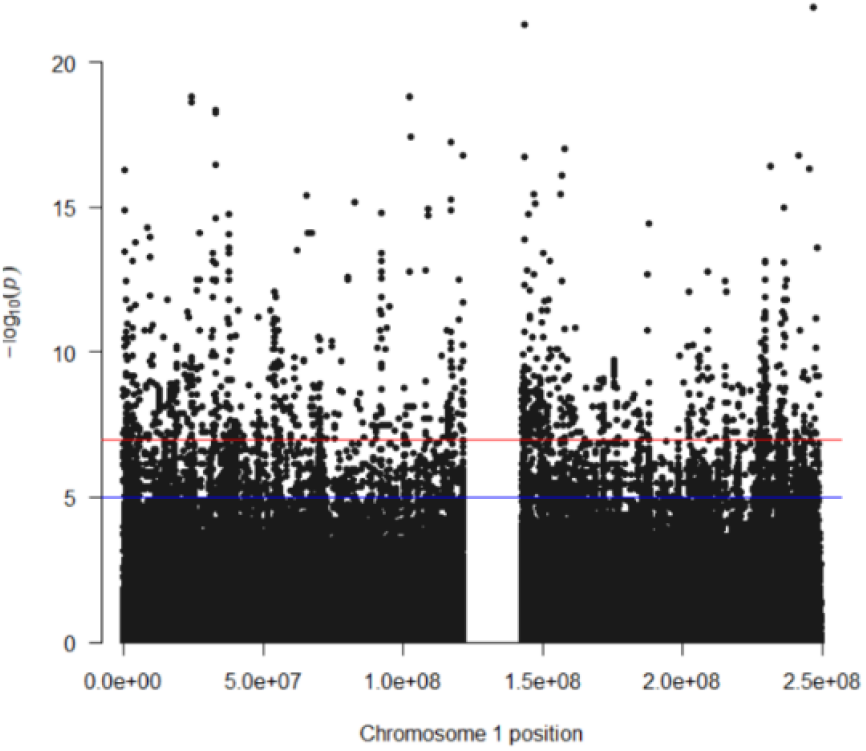
Manhattan Plot of Chromosome 1. Note: Dots above the solid red line were statistically significant.

### 3.2 Chromosome 6

The results of the PCA of the first half of the study, where the HLA genes are located, were almost identical for Japanese, Chinese, and Korean populations, with individual differences more significant than group differences (Fig. 3).

**Fig. 3.**
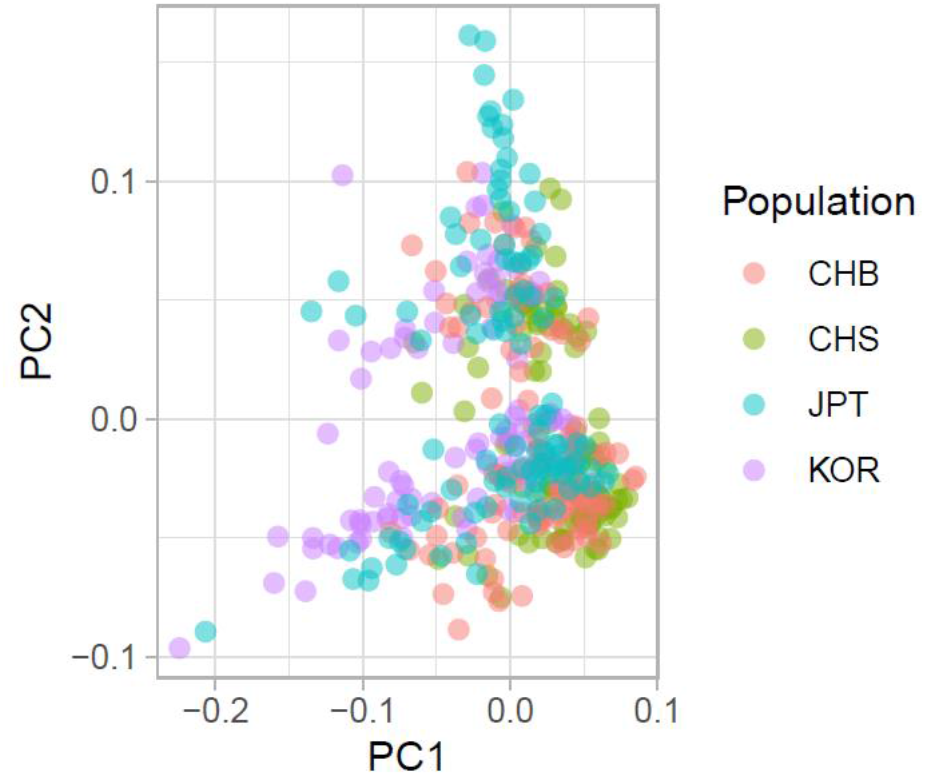
PCA Result of Chromosome 6 (First Half)

The second half of the chromosome showed the distinct characteristics of the Japanese, Chinese, and Korean populations, but the differences were smaller than those at chromosome 1 (Fig. 4).

**Fig. 4.**
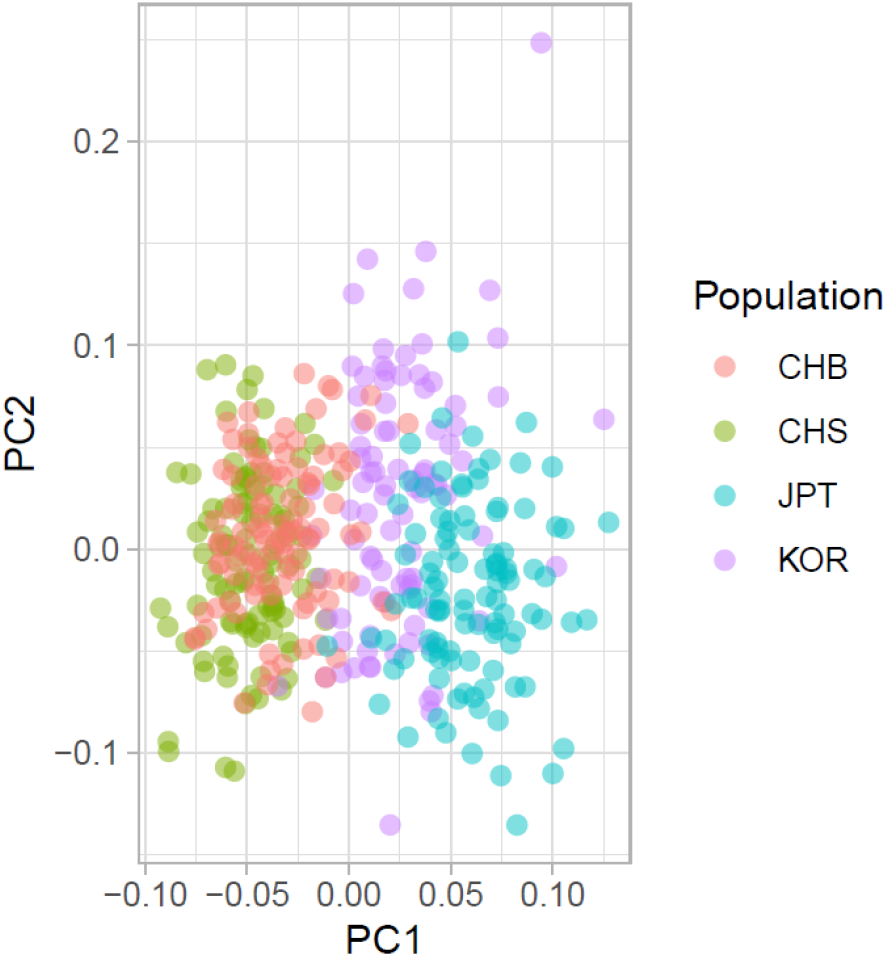
PCA Result of Chromosome 6 (Second Half)

The Manhattan plot at chromosome 6 showed substantial differences in HLA positions, suggesting that the genes of HLA, a major immune system, had been significantly altered and mutated, resulting in extremely large differences between Japanese, Chinese, and Korean groups (Fig. 5).

**Fig. 5.**
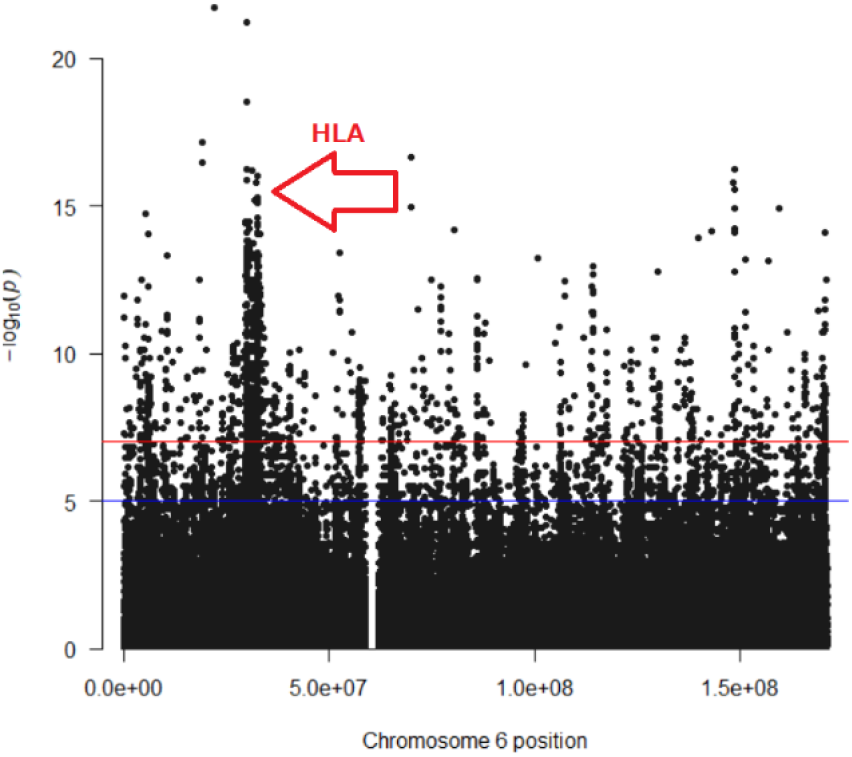
Manhattan Plot of Chromosome 6.

Note: Dots above the solid red line were statistically significant.

### 3.3 Chromosome 9

As with chromosome 1, the Japanese, Chinese, and Korean populations were separated (Fig. 6), although the differences were smaller. Manhattan plot showed no noticeable difference in ABO position (Fig. 7).

**Fig. 6.**
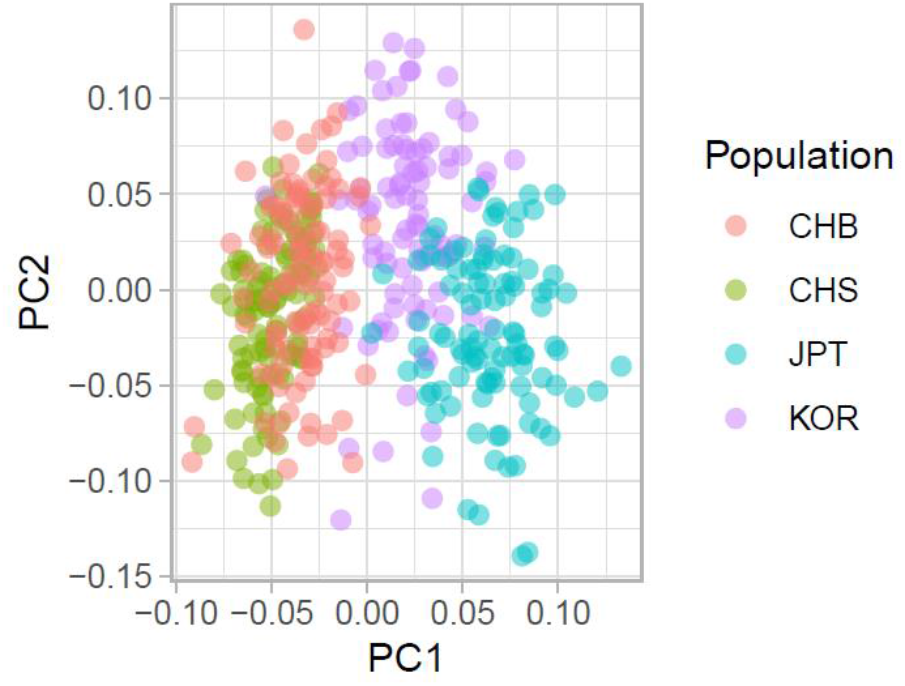
PCA Result of Chromosome 9.

**Fig. 7.**
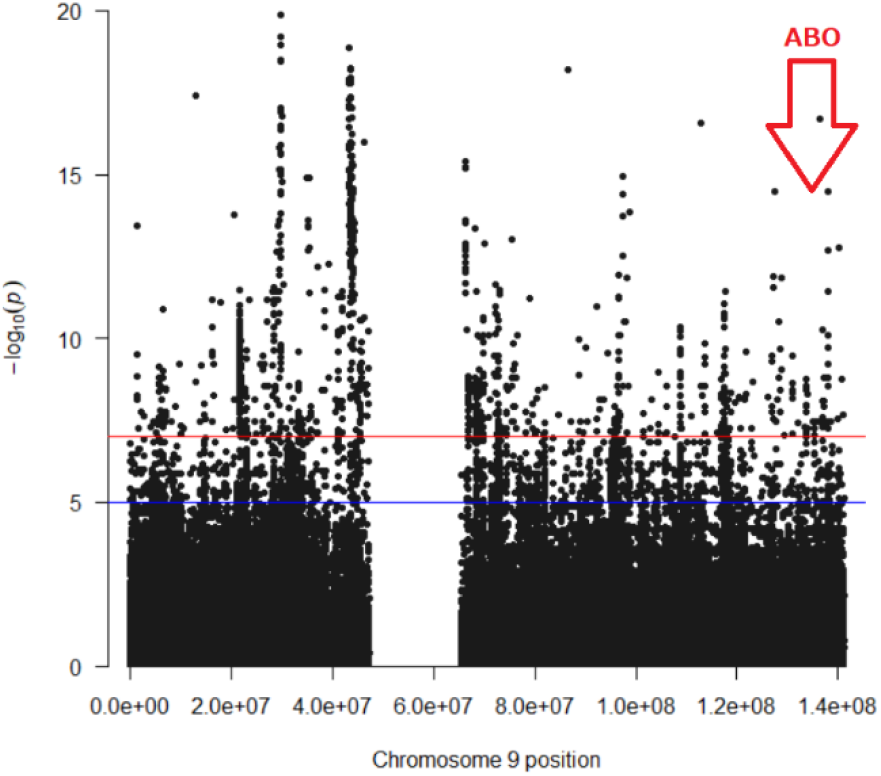
Manhattan Plot of Chromosome 9. Note: Dots above the solid red line were statistically significant.

### 3.4 Chromosome 12

As with chromosome 1, the Japanese, Chinese, and Korean populations were separated. However, the first and second principal component figures showed that three individual subjects had values that were far from their groups (Fig. 8). Manhattan plot showed no noticeable difference in ALDH2 position (Fig.9).

**Fig. 8.**
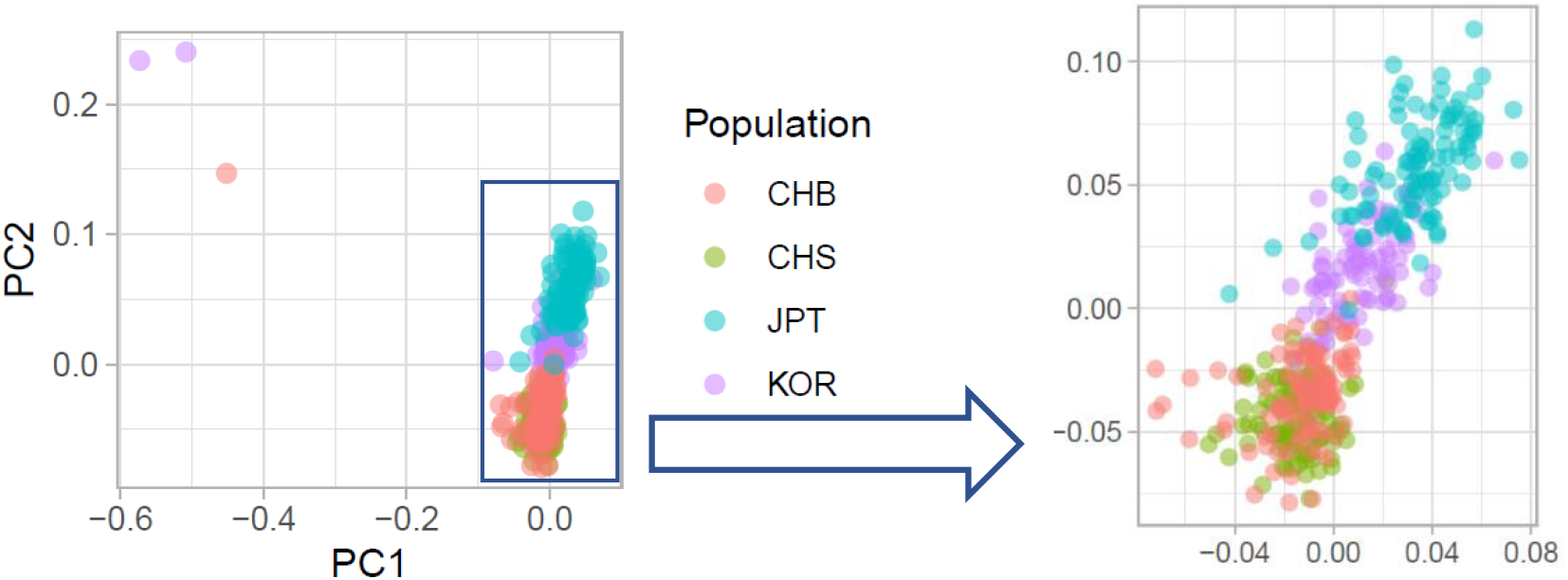
PCA Result of Chromosome 12.

**Fig. 9.**
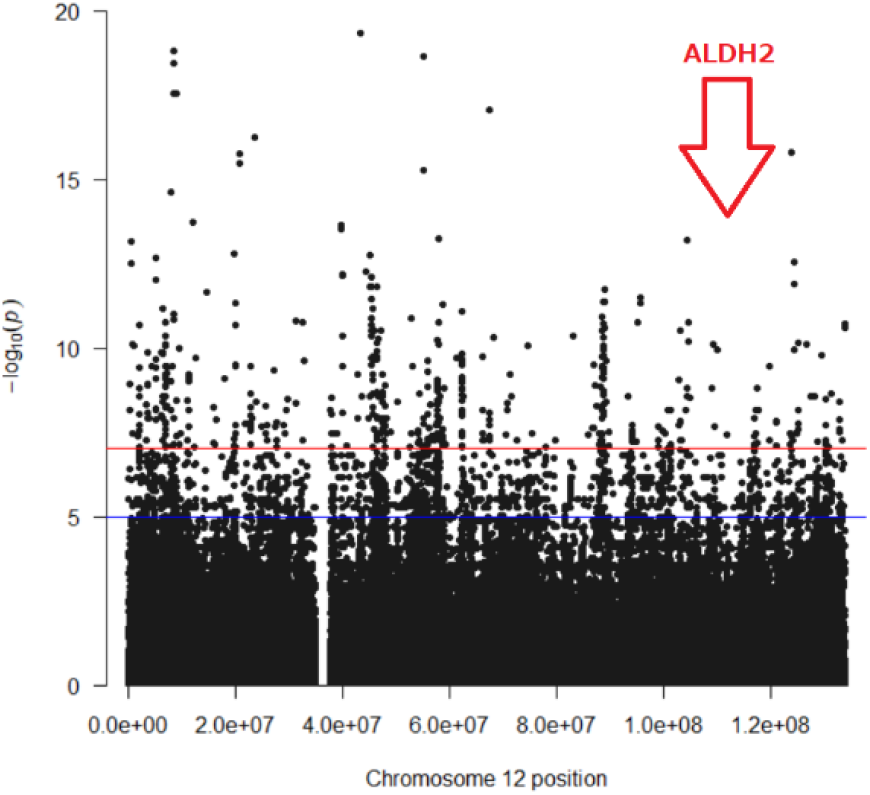
Manhattan Plot of Chromosome 12. Note: Dots above the solid red line were statistically significant.

## 4. Discussion

### 4.1 Estimation of Selection Pressure using Linear Algebra

Looking at the first and second principal components of PCAs, Mahalanobis distances, which indicate differences in human population groups, were considerably smaller in the first half of chromosome 6, which contains HLA, the core of the human immune system where natural selection pressure was strong, in comparison with other chromosomes (Table 1). In the other parts, such as the second half of chromosome 6, the Mahalanobis distance was not very different from that of chromosome 1. These values suggest that natural selection pressure affected the second half of chromosome 6 less than they affected the first half. In general, Mahalanobis distances from JPT (Tokyo, Japan) were also consistent with the physical distances from KOR (Korea), CHB (Beijing, China), and CHS (southern China), in that order.

**Table 1.** Mahalanobis Distances (1st and 2nd Principal Components) from JPT.

| Population | Chromosome Number |  |  |  |  |
| --- | --- | --- | --- | --- | --- |
|  | 1 | 6 (1st Half) | 6 (2nd Half) | 9 | 12 |
| CHB | 6.722 | 0.673 | 3.859 | 4.333 | 5.140 |
| CHS | 7.365 | 0.953 | 4.464 | 4.946 | 5.477 |
| KOR | 5.753 | 0.717 | 1.759 | 2.307 | 2.508 |

As mentioned, the first half of chromosome 6, where the gene for HLA is located, was heavily altered and mutated in SNPs, resulting in extremely large differences among Japanese, Chinese, and Koreans. However, the DNA was very different; the PCA result showed that Japanese, Chinese, and Koreans almost overlapped. This might mean very large individual differences.

### 4.2 Alternative Approach using AI

In a different approach, we used AI, Microsoft Azure Machine Learning, to predict human population groups from the values of the first and the second principal components of PCAs. System parameters were set to defaults. The whole PCA data were divided into two parts, with 70% allocated for training and the remaining 30% for predicting. The result of the AI prediction is shown in Table 2. The larger the Mahalanobis distance (Table 1), the lower the accuracy. This result also suggests that natural selection pressure was strong.

**Table 2.** Result of Human Group Prediction using AI.

|  | Chromosome Number |  |  |  |  |
| --- | --- | --- | --- | --- | --- |
|  | 1 | 6 (1st Half) | 6 (2nd Half) | 9 | 12 |
| Accuracy | 0.967 | 0.733 | 0.895 | 0.950 | 0.918 |
| Algorithm | Gradient Boosting | Random Trees | Logistic Regression | SVM | SVM |

Note: Each accuracy was the maximum value among several algorithms used.

The same dataset and AI were used to analyze the “genomic patterns” of each human population group. Okada et al. [18] suggested that the acetaldehyde decomposition gene (ALDH2) had the strongest natural selection pressure for Japanese people among all genomes. Since human chromosomes are diploid, each SNP has 0 to 2 mutations. Group differences in ALDH2 are shown in Table 3. The higher the number of mutations, the more sensitive to alcohol.

**Table 3.** Number of ALDH2 Mutations by Human Population Group.

| Population | Number of ALDH2 Mutations |  |  |  |
| --- | --- | --- | --- | --- |
|  | 0 | 1 | 2 | Total |
| JPT | 59<br>(56.7%) | 40<br>(38.5%) | 5<br>(4.8%) | 104<br>(100%) |
| CHB | 74<br>(71.8%) | 25<br>(24.3%) | 4<br>(3.9%) | 103<br>(100%) |
| CHS | 57<br>(54.3%) | 39<br>(37.1%) | 9<br>(8.6%) | 105<br>(100%) |
| KOR | 66<br>(72.5%) | 23<br>(25.3%) | 2<br>(2.2%) | 91<br>(100%) |

We predicted human groups based on the pattern of mutation counts for all 100 SNPs, including ALDH2 and those before and after (POS 112211833 - 112263268). Similarly, the whole data were divided into two parts, with 70% allocated for training and the remaining 30% for predicting. The accuracy was 63.9% (algorithm: XG Boost Classifier), more than twice the 25% probability of a chance match to one of the four population groups (JPT, CHB, CHS and KOR). The result also suggests that natural selection pressure was strong.

### 4.3 Effect of Selection Pressure

From the results above, it can be inferred that natural selection by infectious diseases does not have a “positive” effect on specific genes, but rather a “negative” effect. In other words, genes change in the direction of larger diversity. This implies that individuals with genes that make them more susceptible to infectious diseases are more likely to go extinct, as there is a greater likelihood that they will decline rapidly since they will not be able to pass their genes on to the next generation. The large genetic variation probably means that it is advantageous for the survival of the species for the immune system to diversify, rather than the “survival of the fittest.” This is similar to the situation with resistant bacteria or COVID-19; the latter is still mutating to escape vaccines.

It is estimated that Japan, being an island nation, has a relatively small population influx from outside compared to continental nations such as China and Korea. However, in light of the above, it may not always be a meaningful comparison to consider how much of the ancient Japanese (Jomon) genes are inherited by modern Japanese people. Japan has been separated from the Asian continent since about 20,000 years ago, and from that point until about 3,000 years ago (the Jomon period), Japan was not an agricultural society. Later (the Yayoi period), with the arrival of paddy rice cultivation, the society transformed into an agricultural one, the staple food changed to rice, intense infectious diseases such as tuberculosis became prevalent, and the environment changed drastically [21]. Therefore, it is assumed that among the Jomon people, those individuals who could not cope with such changes died without children. Therefore, it is highly likely that the modern Japanese people who have the genes to cope with infectious diseases are considerably different from those of the Jomon people, who did not live in an agricultural society until 3,000 years ago.

### 4.4 Influence of Paddy Rice Cultivation

The “low tolerance for alcohol” gene, which is said to have originated in the Yangtze River region, may be another example [20]. More than 6,000 years ago, many people began to gather and live near the Yangtze River floodplain, which was suitable for rice cultivation. At that time, because of a poor sanitary environment, food was often contaminated with harmful microbes and other substances that could cause infectious diseases. At such a time, alcoholic beverages made from rice were thought to be useful.

When people with a low tolerance for alcohol, or weak acetaldehyde decomposition gene (ALDH2), drank alcoholic beverages, the level of acetaldehyde, a highly poisonous substance that cannot be decomposed, would increase in his or her body. However, it appears that the poison might have also served as a drug that attacked harmful microbes. On the other hand, people without this weak gene had less acetaldehyde in their bodies and could not suppress those harmful microorganisms. Thus, people with the “low tolerance for alcohol” gene were more likely to survive and overcome infectious diseases.

In other words, it is possible that people in rice paddy farming areas felt selection pressure to develop a low tolerance for alcohol in order to protect themselves from infectious diseases. This “low tolerance for alcohol” gene, ALDH2, was eventually introduced to the Japanese archipelago along with the rice culture; over 40% of the present Japanese population has this gene. It is also thought that many ancient Japanese before rice paddy cultivation (Jomon people) did not have this gene.

Tuberculosis (TB) is another infectious disease thought to have arrived in Japan along with rice paddy cultivation [21]. People with blood types B and AB have the same type B antigen as Mycobacterium Tuberculosis, making it difficult for their immune system to function and making them susceptible to infection [23]. On the other hand, people with blood types A and O, which do not carry the type B antigen, are less susceptible to TB infection. In East Asia, paddy rice cultivation is prevalent in Japan, southern China, and Korea, where types A and O tend to be more common than types B and AB [24]. However, due to improved sanitary conditions in today’s East Asia, it would be difficult to substantiate the above hypotheses.

### 4.5 Interpretation of PCA and Manhattan Plot

According to the PCA and the Manhattan plot for each chromosome conducted in this study, the results seem to vary considerably depending on the selection pressure, human group selection methods, and sample sizes.

For example, on chromosome 6, which contains the HLA gene, the Mahalanobis distances for Japanese, Chinese, and Koreans are relatively close compared to other chromosomes according to PCA analysis (Table 1). However, Manhattan plot analysis results show that the differences between Japanese and Chinese or Japanese and Korean SNPs are larger than the other chromosomes (Fig. 5), which were the exact opposite.

In addition, ALDH2, the “low tolerance for alcohol” gene commonly found in Japan, southern China, and Korea, is said to differ more from other human groups [15], but significant differences were not found in PCA or Manhattan plot (Figs. 8-9), whereas differences were found in AI-based analysis (Table 3).

In light of the above, when making comparisons among human groups, sufficient attention should be paid not only to sample selection methods and sample sizes, but also to the linear algebraic nature of PCA and Manhattan plot. A comprehensive perspective will also be required when interpreting the results.

## 5. Conclusion

GWAS is useful for comparing characteristics of human groups. However, genomes may change rapidly over time when selection pressures, such as environmental changes, are strong. In particular, the immune system seems to be very sensitive to environmental changes. Therefore, there will be a need to perform GWAS not only on whole genomes, but also at the level of individual chromosomes when necessary.

Japanese, Chinese, and Korean people form distinctly different groups genetically. However, the sample size for this study is small (403 individuals), the targeted samples are limited to the East Asian population, and only a few basic methods were used for data analysis. Studies with a larger global dataset and methodological innovations will be needed to get one step closer to the truth.

## Reference

1. 2003: Human Genome Project Completed. National Human Genome Research Institute. Retrieved from https://www.genome.gov/25520492/online-education-kit-2003-human-genome-project-completed

2. https://www.internationalgenome.org

3. 1000 Genomes Project Consortium, Auton A, Brooks LD, Durbin RM, Garrison EP, Kang HM, Korbel JO, Marchini JL, McCarthy S, McVean GA, Abecasis GR. A global reference for human genetic variation. Nature. 2015; 526(7571): 68–74. DOI: 10.1038/nature15393.

4. Jeon S, Bhak Y, Choi Y, Jeon Y, Kim S, Jang J, Jang J, Blazyte A, Kim C, Kim Y, Shim J, Kim N, Kim YJ, Park SG, Kim J, Cho YS, Park Y, Kim HM, Kim BC, Park NH, Shin ES, Kim BC, Bolser D, Manica A, Edwards JS, Church G, Lee S, Bhak J. Korean Genome Project: 1094 Korean personal genomes with clinical information. Sci Adv. 2020; 6(22): eaaz7835. DOI: 10.1126/sciadv.aaz7835.

5. Bai H, Guo X, Narisu N, Lan T, Wu Q, Xing Y, Zhang Y, Bond SR, Pei Z, Zhang Y, Zhang D, Jirimutu J, Zhang D, Yang X, Morigenbatu M, Zhang L, Ding B, Guan B, Cao J, Lu H, Liu Y, Li W, Dang N, Jiang M, Wang S, Xu H, Wang D, Liu C, Luo X, Gao Y, Li X, Wu Z, Yang L, Meng F, Ning X, Hashenqimuge H, Wu K, Wang B, Suyalatu S, Liu Y, Ye C, Wu H, Leppälä K, Li L, Fang L, Chen Y, Xu W, Li T, Liu X, Xu X, Gignoux CR, Yang H, Brody LC, Wang J, Kristiansen K, Burenbatu B, Zhou H, Yin Y. Whole-genome sequencing of 175 Mongolians uncovers population-specific genetic architecture and gene flow throughout North and East Asia.N at Genet. 2018; 50(12):1696–1704. DOI: 10.1038/s41588-018-0250-5.

6. Tadaka S, Katsuoka F, Ueki M, Kojima K, Makino S, Saito S, Otsuki A, Gocho C, Sakurai-Yageta M, Danjoh I, Motoike IN, Yamaguchi-Kabata Y, Shirota M, Koshiba S, Nagasaki M, Minegishi N, Hozawa A, Kuriyama S, Shimizu A, Yasuda J, Fuse N; Tohoku Medical Megabank Project Study Group, Tamiya G, Yamamoto M, Kinoshita K. 3.5KJPNv2: an allele frequency panel of 3552 Japanese individuals including the X chromosome. Hum Genome Var. 2019; 6:28. DOI: 10.1038/s41439-019-0059-5.

7. Reich D. Who We Are and How We Got Here: Ancient DNA and the new science of the human past. 2019; Oxford: Oxford University Press.

8. Cao Y, Li L, Xu M, Feng Z, Sun X, Lu J, Xu Y, Du P, Wang T, Hu R, Ye Z, Shi L, Tang X, Yan L, Gao Z, Chen G, Zhang Y, Chen L, Ning G, Bi Y, Wang W; ChinaMAP Consortium. The ChinaMAP analytics of deep whole genome sequences in 10,588 individuals. Cell Res. 2020 Sep;30(9):717–731. DOI: 10.1038/s41422-020-0322-9.

9. Yoo SK, Kim CU, Kim HL, Kim S, Shin JY, Kim N, Yang JSW, Lo KW, Cho B, Matsuda F, Schuster SC, Kim C, Kim JI, Seo JS. NARD: whole-genome reference panel of 1779 Northeast Asians improves imputation accuracy of rare and low-frequency variants. Genome Med. 2019; 11(1): 64. DOI: 10.1186/s13073-019-0677-z.

10. Robbeets M, Bouckaert R, Conte M, Savelyev A, Li T, An DI, Shinoda KI, Cui Y, Kawashima T, Kim G, Uchiyama J, Dolińska J, Oskolskaya S, Yamano KY, Seguchi N, Tomita H, Takamiya H, Kanzawa-Kiriyama H, Oota H, Ishida H, Kimura R, Sato T, Kim JH, Deng B, Bjørn R, Rhee S, Ahn KD, Gruntov I, Mazo O, Bentley JR, Fernandes R, Roberts P, Bausch IR, Gilaizeau L, Yoneda M, Kugai M, Bianco RA, Zhang F, Himmel M, Hudson MJ, Ning C. Triangulation supports agricultural spread of the Transeurasian languages. Nature. 2021; 599(7886): 616–621. DOI: 10.1038/s41586-021-04108-8.

11. Jinam TA, Kanzawa-Kiriyama H, Saitou N. Human genetic diversity in the Japanese Archipelago: dual sructure and beyond. Genes Genet Syst. 2015;90(3):147–52. DOI: 10.1266/ggs.90.147.

12. Cooke NP, Mattiangeli V, Cassidy LM, Okazaki K, Stokes CA, Onbe S, Hatakeyama S, Machida K, Kasai K, Tomioka N, Matsumoto A, Ito M, Kojima Y, Bradley DG, Gakuhari T, Nakagome S. Ancient genomics reveals tripartite origins of Japanese populations. Sci Adv. 2021; 7(38): eabh2419. DOI: 10.1126/sciadv.abh2419.

13. Gelabert P, Blazyte A, Chang YJ, Fernandes DM, Jeon SW, Hong JG, Yoon JY, Ko YM, Oberreiter V, Cheronet O, Özdoğan KT, Sawyer S, Yang SH, Greytak EM, Choi HS, Kim JG, Kim JI, Bae KD, Bhak J, Pinhasi R. Diverse northern Asian and Jomon-related genetic structure discovered among socially complex Three Kingdoms period Gaya region Koreans. BioRxiv. 2021.10.23.465563 DOI: 10.1101/2021.10.23.465563

14. ftp://koref.biodisk.org/

15. Genome Map, Ministry of Education, Culture, Sports, Science and Technology Japan. https://www.mext.go.jp/stw/common/pdf/series/genome_map/genome_English_2nd_A3.pdf

16. Hoiland RL, Fergusson NA, Mitra AR, Griesdale DEG, Devine DV, Stukas S, Cooper J, Thiara S, Foster D, Chen LYC, Lee AYY, Conway EM, Wellington CL, Sekhon MS. The association of ABO blood group with indices of disease severity and multiorgan dysfunction in COVID-19. Blood Advances. 2020; 4(20): 4981–4989. DOI: 10.1182/bloodadvances.2020002623

17. Severe Covid-19 GWAS Group. Genomewide Association Study of Severe Covid-19 with Respiratory Failure. New England Journal of Medicine. 2020; 383(16): 1522–1534. DOI: 10.1056/NEJMoa2020283

18. Okada Y, Momozawa Y, Sakaue S, Kanai M, Ishigaki K, Akiyama M, Kishikawa T, Arai Y, Sasaki T, Kosaki K, Suematsu M, Matsuda K, Yamamoto K, Kubo M, Hirose N, Kamatani Y. Deep whole-genome sequencing reveals recent selection signatures linked to evolution and disease risk of Japanese. Nat Commun. 2018; 9(1): 1631. DOI: 10.1038/s41467-018-03274-0.

19. https://www.cog-genomics.org/plink/

20. https://cran.r-project.org/

21. Okazaki K, Takamuku H, Yonemoto S, Itahashi Y, Gakuhari T, Yoneda M, Chen J. A paleopathological approach to early human adaptation for wet-rice agriculture: The first case of Neolithic spinal tuberculosis at the Yangtze River Delta of China. International Journal of Paleopathology. 2019; 24: 236–244. DOI: 10.1016/j.ijpp.2019.01.002

22. NHK (Japan Broadcasting Corporation). Evolutionary fate makes you want to drink: the unknown truth about alcoholic beverages. Retrieved from https://www.nhk.or.jp/special/plus/articles/20200127/index.html

23. Oike Y, Kikuchi Y, Kushibiki H, Kudo M, Kobori T, Araya K. The Relationship between the ABO Blood-Groups and Tuberculosis. Nihon Naika Gakkai Zasshi. 1954; 42(11): 835–838. DOI: 10.2169/naika.42.11_835

24. A.E. Mourant Ae, Kopec AC, Domaniewska-Sobczak K. The Distribution of the Human Blood Groups and Other Polymorphisms (Monographs on Medical Genetics). 1976; Oxford: Oxford University Press.

